# Vocabulary Matters: An Annotation Pipeline and Two Deep Learning Algorithms for Enzyme Named Entity Recognition

**DOI:** 10.1101/2023.06.23.546229

**Authors:** Meiqi Wang, Avish Vijayaraghavan, Tim Beck, Joram M. Posma

## Abstract

Enzymes are indispensable substances in many biological processes. With biomedical literature growing exponentially, it becomes more difficult to review the literature effectively. Hence, text-mining techniques are needed to facilitate and speed up literature review. The aims of this study are to create a corpus with annotated enzymes to train and evaluate enzyme named-entity recognition (NER) models. A novel pipeline was built using a combination of dictionary matching and rule-based keyword searching to automatically annotate enzyme entities in over 4,800 biomedical full texts. Two Bidirectional Long Short-Term Memory (BiLSTM) networks using BioBERT and SciBERT as tokeniser and word embedding layers were trained on this corpus and evaluated on a manually annotated test set of 526 full-text publications. The dictionary- and rule-based annotation pipeline achieved an F1-score of 0.863 (precision 0.996, recall 0.762). The SciBERT-BiLSTM model (F1-score 0.965, precision 0.981, recall 0.954) largely out-performed the BioBERT-BiLSTM model (F1-score 0.955, precision 0.981, recall 0.937). This study contributed a novel dictionary- and rule-based automatic pipeline with almost perfect precision which runs in a matter of seconds on a standard laptop. Both deep learning (DL) models achieved state-of-the-art performance (F1*>*0.95) for enzyme NER, with the SciBERT-based model outperforming the BioBERT-based model in terms of recall, demonstrating the vocabulary used by models matters. The proposed pipeline with the DL models can facilitate more effective enzyme text-mining and information extraction research for literature review and are the first algorithms specifically for enzyme NER.

**Availability:** All codes are available for automatic annotation and model training (including data), with instructions on how to deploy the model on new text, from https://github.com/omicsNLP/enzymeNER.

## Introduction

Enzymes are proteins that regulate biochemical reactions by decreasing the activation energy required for the chemical conversion from a substrate into a product to occur. The largest online enzyme database, BRENDA (1), contains over 100K different enzyme names that are obtained by manual extraction from literature, semi-automatic text mining or by integration from other sources and is growing exponentially (2). Manual extraction of information from large numbers of publications is time-consuming, labour-intensive and may lead to human errors. With the biomedical literature expanding at an increased rate (3), there is a clear need to develop new methods to automatically identify and extract enzyme names from the biomedical scientific literature.

Natural language processing (NLP) is a sub-field of artificial intelligence that uses statistics-based techniques to understand human language, and has multiple different applications such as text prediction, classification, summarisation and translation, as well as speech recognition (4). Recent advances in the field have resulted in obtaining high performance of various NLP tasks with deep learning (DL) models and domain-specific word embeddings. Named-entity recognition (NER) is an NLP task that locates and identifies specific entities in texts and classifies them into predefined categories. In recent years, NER models using DL models have flourished given their high performance on many tasks (5), using DL architectures such as bidirectional long-short term memory (BiLSTM) based recurrent neural networks (6, 7) and transformers (8–11). There has been much recent success of biomedical natural language processing (bioNLP) such as for gene/protein, disease and chemical entity recognition with fine-tuned pre-trained language models (12–15), as well as for reading clinical notes in electronic health records (16).

The concept of pre-training refers to initialising network weights based on a large-scale corpus (a collection of texts/sentences). This is done to generate distributed representations of words, or N-dimensional vectors, as inputs for subsequent NER models. In NLP, this processing is known as word embedding which is a type of pre-trained language model. The word embedding process can not only yield machine-understandable expressions of real words, but also capture semantic and syntactic properties of a word which might not be explicitly present without word embedding (5), and this is context- and domain-dependent. Until a few years ago, static word embedding such as Word2Vec (17) and GloVe (18) were predominantly used. These have now been replaced by contextualised word embeddings that enhance the DL model performance (19) by focusing on neighbouring words. These pre-trained models can then be finetuned (changing network weights) for different downstream NLP tasks using transfer learning (5). The choice of pretrained model has a big influence on the final performance because general domain datasets, such as those to train Bidirectional Encoder Representations from Transformers (BERT) (9) (trained on Wikipedia and Google Books corpus), may not contain the vocabulary and word embeddings that are necessary to process text from biomedical literature (19). BioBERT is a biomedical language representation model that was pre-trained on biomedical domain corpora (PubMed abstracts with 4.5B words and PubMed Central (PMC) full-text publications with 13.5B words) after initialisation with BERT-weights and can be fine-tuned for NER, relation extraction and question answering text-mining problems and has shown state-of-the art performance for several NER tasks (12). SciBERT (11) is a related model which was initialised from scratch and trained on full texts publications from the Semantic Scholar platform (20), and thus does not inherit any vocabulary from BERT making it potentially more useful for bioNLP tasks as it was trained on scientific publications from the biomedical domain (82%) and computer science (18%). Training of models for biomedical NER relies on the availability of biomedical text corpora in machine-readable format (21). Most corpora used for training (and evaluating) bioNLP algorithms are those either curated by the BioCreative committee or NCBI team and contain annotations for different entities including for diseases (22, 23), genes/proteins (24– 26), mutations (27) and chemical compounds (23, 28, 29). The availability of corpora dictates the developments made in the field of bioNLP, with mostly incremental gains over prior methods. For the task of enzyme NER no corpus exists that is specific for this task, thus only corpora with annotated protein entities can be used. However the specificity is a problem since not all proteins are enzymes, thereby training on protein corpora for enzyme recognition will inflate false positives. Moreover, the existing corpora with protein (and other) annotations come mostly from abstracts, as can be seen from data used by CollaboNet(30). In this article we aim to fill this gap by describing our dictionary-based annotation workflow to create an annotated enzyme corpus in machine-readable format, curated from full-text publications. We then describe two DL-based NER models trained using different word embeddings and evaluate these on a manually annotated test set.

## Materials and Methods

### Data - Publications

The corpus was created from PMC Open Access publications from four different omics domains: genome-wide association studies (GWAS), proteomics, metabolomics and microbiome. The GWAS and metabolomics publications that were used we described previously (31, 32), the proteomics and microbiome publications were acquired in a similar manner as the metabolomics publication (32). For proteomics the field keyword search terms were ‘proteomics’, ‘proteome’, ‘proteomic’ and ‘proteome-wide’, the sample keyword search terms were ‘urine’, ‘urinary’, ‘blood’, ‘serum’, ‘plasma’, ‘faecal’, ‘faeces’, ‘fecal’, ‘feces’, ‘stool’, ‘cerebrospinal fluid’, ‘CSF’ and ‘biofluid’, with similar disease search terms. For the microbiome corpus OA publications relating to airway, faecal, skin, urinary and vaginal microbiomes were searched for with additional keywords such as ‘16S’, ‘rRNA’, ‘sequencing’, ‘shotgun’, ‘Illumina’, ‘MiSeq’ and ‘PCR’.

The publications were converted from HTML, standardised and converted to the machine-readable BioC-JSON format using Auto-CORPus (31). First, Auto-CORPus converts the main text of each publication from HTML to BioC JSON format. In this file, the pipeline splits the publications into different sections using the Information Artifact Ontology (IAO) (33) and in each section, each paragraph forms a single ‘text’-feature. Second, Auto-CORPus transforms tables inside publications to a table-JSON format, and it extracts abbreviations from both the main text and from separate abbreviations sections (if available) within the main text and generates another JSON file for abbreviations with linked full definitions. Here, we use the main text (BioC-JSON) and abbreviation JSON files for pre-processing and to build the datasets for DL models.

### Data - Enzyme Nomenclature

A list of enzymes codes with names and synonyms was downloaded on 28/10/2021 using the KEGG REST API (https://www.kegg.jp/kegg/rest/keggapi.html). This list contains over 28K enzyme names/synonyms that were used for the dictionary matching here. Several changes were made to this dictionary to improve the matching step later as the matching list in this study. Missing entities were added that have the same meaning but different spelling styles compared with other items within the list. For example, some enzymes connect different words using hyphens while others use spaces instead, whereas others use Greek letters opposed to spelling these out using the Latin alphabet. Therefore, additional entries were created in the dictionary to account for these differences and capture more entities when matching the texts.

### Annotation Pipeline

The second improvement made relates to speeding up the searching process. We reconstruct the dictionary list in a novel hierarchical format by clustering entities with the same features into smaller streams based on enzyme classes. We performed this based on analysis of the enzyme nomenclature, where most (but not all) enzyme names end in ‘ase’. We use the seven categories of enzymes based on the different chemical reactions they are involved in. For example, enzymes involved in oxidation reactions are likely ending in ‘oxidase’ whereas ‘reductase’ enzymes perform reduction reactions. Using this structure, we classify the items within the list in a tree-structured (hierarchical) dictionary, in that way, when searching through the text, the pipeline can use regular expression (RegEx) rules to search the node-words from the root of the word to determine which branch-list to use to speed up the searching and auto-annotation process.

Supervised NER algorithms rely on annotated corpora, however none exist for enzymes. We created an enzyme-specific corpus using a fast, automated pipeline for annotation using the hierarchical dictionary of enzyme terms. The pipeline uses rule-based RegEx to assist with the search process of potential entities in full-text articles. The BioC JSON files from Auto-CORPus were used as input to the annotation pipeline, with the enzyme entities in each paragraph added to the ‘annotations’ section of the JSON document which includes a unique id, an external identifier (EC number) if a direct match is found (if no direct match is found this is replaced by the root word) and the location in the text (based on the offset of the paragraph in the entire article).

The pipeline consists of two search steps to find entities. First, direct matches of dictionary terms in the text (exact match to a known entity). Second, using keyword searching using root enzyme terms (e.g. ‘transferase’) to perform rule-based fuzzy matching to identify entities that do not directly match to elements from the dictionary. Specifically, the pipeline utilizes the spaCy package (v3.2) to split each paragraph into individual sentences, in which each word is contrasted with patterns created by the searching rules based on the structure of enzyme dictionary.

If a (root) word is matched, the pipeline will then first search throughout the given node in the list for a content-matched item (first step). In this step, the greedy algorithm was used to ensure the matched entity is the one with the maximum length in this sentence. For example, the term ‘aliphatic alcohol dehydrogenase’ also contains other matches to terms ‘alcohol dehydrogenase’ and ‘dehydrogenase’, and this approach ensures any terms contained within another are not selected.

If no direct match is found, the pipeline continues with step 2. The principle of partial searching is that if a word is regarded as a part of an enzyme name, such as a word that is matched with a node-word in the dictionary and words ending with ‘ase’ are considered as a part of a potential enzyme. The next steps considers the words preceding and following the identified entity and evaluates if these are part of the enzyme entity. In this step common words ending in ‘-ase’, e.g., ‘phase’ and ‘database’, are ignored. To establish the number of words before and after an entity to consider, we analysed the known enzyme names in the dictionary. Over 92% enzyme entities in the list end in ‘-ase’, and from these over 99% consist of five words or less, indicating that we can restrict our search to four words before the identified ‘ase’-word. For words after the entity we use additional RegEx rules as most of the contents after the ‘ase’-word are enclosed in brackets, but cases exist where there is a single number or Greek letter. Therefore, we created rules based on these features to support backward searching and adding these to the identified entity.

To increase the number of training cases, we also include sentences in which abbreviations of enzymes appear. We applied the pipeline to the list of definitions (abbreviations output JSON file from Auto-CORPus) in the article, and the abbreviations of any enzymes identified are then searched in the main text. These abbreviations are annotated in the same way as full terms, however for training these are replaced by their full definitions. Abbreviations are not used by themselves as this can increase the number of false positives as some abbreviations can have different definitions in different articles.

### Training, Validation, Test Splits

The corpus of 3,525 full-text publications that contained any enzyme mention was split 75:10:15 into training:validation:test sets with each portion containing an equal proportion of GWAS, proteomics, metabolomics and microbiome articles (Table 1). The training and validation sets were used to construct the deep learning models, with the test set separately under-going manual annotation and used to evaluate the performance of the deep learning algorithms. The training and validation sets were annotated using the annotation pipeline.

**Table 1.** Summary of the enzyme corpus in terms of total (n) publications (pubs), and the splitting into training (train), validation (val) and test sets.

| Domain | Pubs (n) | Train (n) | Val (n) | Test (n) |
| --- | --- | --- | --- | --- |
| GWAS | 884 | 664 | 88 | 132 |
| proteomics | 446 | 336 | 44 | 66 |
| metabolomics | 1,006 | 756 | 100 | 150 |
| microbiome | 1,189 | 893 | 118 | 178 |

Only the sentences containing enzyme entities constitute the dataset for training models. Therefore, we extract those annotated sentences to build up the final corpus for DL models. Each sentence gets a unique ID which consists of the PM-CID (i.e., paper ID with PMC identifier), a sentenceID that determines which section (e.g., Results) and paragraph the sentence belongs to. Unlike our metabolite NER model (32) where abbreviations of metabolites were kept in, here we replaced all abbreviations in the full text with the definitions from the abbreviation output files. To increase the number of annotations for training, we consider not only the textual abstract, methods, results, and discussion sections (as done for metabolite NER), but we also include the introduction and references sections as supplement.

The test set consists of 526 full-text publications. These were annotated in a three-stage process. First, the articles were annotated using the automatic annotation pipeline (see below). Second, two annotators independently annotated the 526 articles using TeamTat (34), resolved incorrect annotations from the pipeline, added missing concept identifiers (EC numbers) and annotated missing entities. The final step was done in collaborative mode in TeamTat where all annotators can see the annotations made by both annotators in the independent mode. TeamTat indicates any conflicts between annotators that need to be resolved by the annotators (and mediated by a third annotator in case of disagreements). Conflicts include cases where only one annotator has annotated an entity, where annotated entities are overlapping and where the concept ID is different between annotators.

### Deep Learning Models

In this study, the BiLSTM network structure is used because of its state-of-the-art performance achieved in NER tasks in the chemical domain. ChemListem (35), a type of BiLSTM model, outperformed other methods for chemical entities recognition and was validated by the CEMP BioCreative V.5 challenge (36), and MetaboListem and TABoLiSTM (32) achieved similar performance for metabolite named entity recognition. To further optimise ChemListem, the ability to accurately capture the features of enzyme entities can be enhanced by replacing GloVe embedding method with other word embedding models which have displayed accurate results in biomedical text mining.

Here, we use two different word embeddings which are BioBERT (12) and SciBERT (11), and then separately combine the BiLSTM architecture from ChemListem to improve the performance of the enzyme NER task. As our previous study (32) demonstrated, BioBERT embedding outperforms BERT, which is reasonable considering that BioBERT was pre-trained on a large sum of publications from biomedical domain. Therefore, we use BioBERT instead of BERT as word embedding to compare the performance with SciBERT. All these models are trained on the dataset we created with the annotation pipeline and are evaluated in the gold-standard test set (i.e., manually annotated dataset).

### Pre-processing, Tokenisation and Word Embedding

Prior work used a Python translation of the Open-Source Chemistry Analysis Routines (OSCAR) software (37) as tokeniser (32, 35). In this study, BioBERT (12) (biobert-base-cased-v1.2) and SciBERT (11) (scibert_scivocab_cased) are employed as transformer-based word tokenisers to generate the vocabulary and corresponding word embedding architectures used in the different NER models.

The SOBIE system is applied to mark each token as a specific tag, in which ‘O’ (for outside) labels a token that is not part of an entity, ‘S’ refers to singleton that marks a one-word token (i.e., the whole of an entity), ‘B’ marks the beginning word of a multi-word token, ‘I’ marks one inside an entity, and ‘E’ marks one at the end. A pre-classifier subsystem was implemented as in prior work (32, 35) using a random forest. The pre-classifier works by generating name-internal binary features that are either ‘O’ or ‘SBIE’ labels and assign a probability to each token for it to be part of an enzyme name (S, B, I or E). These tokens are segmented by BioBERT or SciBERT, respectively, to match the corresponding models. Afterwards, these pre-trained features are passed to the BiLSTM network.

### BiLSTM Network

The BiLSTM model contains two inputs that are concatenated; the first input layer is a Conv1D with a dropout of 0.5 to process the name-internal features from the pre-classifier, this is a convolutional layer with 256 neural dimensions. The second input are the features from the word embedding layer (here BioBERT and SciBERT are separately applied to this layer) where text tokens are converted to a sequence of integers. These contextual embeddings were initialised by pre-trained weights of the transformer models with a 768 hidden dimensional size, feeding the output from the embedding model through a SpatialDropout1D layer with a dropout rate of 0.1 to prevent overfitting. Both input layers are concatenated to result in a 1,024 dimensional vector (256+768) as final input into the BiLSTM network that contains a 64-dimensional output space. The activation function selection is with a tanh (hyperbolic tangent), with a sigmoid activation function applied to the recurrent step of the network. The regularisation penalty used for the kernel weights matrix was l1_l2 in which both the l1 and l2 loss are limited up to 1*×*10^-6^. The output layer is a time-distributed dense layer with a softmax activation function to ensure the sum of the 5 output probabilities (‘O’, ‘S’, ‘B’, ‘I’, and ‘E’) for each token totals to 1.

Models were trained for 10 epochs with each epoch trained in mini batches on a workstation with an Intel Xeon W-2125 CPU with 128GB RAM (4×32GB) and an NVIDIA RTX5000 GPU with 16GB memory and 3,072 CUDA parallel-processing cores. The model with the highest F1 score (see below) for the validation dataset is stored and used to evaluate on the manually annotated test set.

### Comparison of our model on protein NER using BERN2

We compare our enzyme NER results with the BERN2 multi-task model (15), which is based on BioBERT, for the gene/protein entity tag as no enzyme-specific NER model currently exists. The BERN2 model can be accessed remotely through a web API (http://bern2.korea.ac.kr/) but this is limited to 300 requests per 100 seconds per user. For a quicker turnaround time on our large corpus, we followed the authors’ guidelines in the BERN2 repository (https://github.com/dmis-lab/BERN2) to install and run a local version of the model on a Linux workstation with an Intel Core i7 CPU (10700K, 3.8GHz, 8 Core) with 32GB RAM and NVIDIA GeForce RTX 3080 GPU with 10GB memory. Some slight modifications were made to set up the BERN2 model on a Linux workstation with a GPU. In the “run_bern2.sh” script, we changed “python” to “python3”. Every time it runs, BERN2 creates a “log/” directory to record run details which can help identify the location of a problem. From this, we found an issue with the installation of `GNormPlusJava` (the gene/protein normalisation tool) which required us to redownload an alternate version of the CRF++ tool. BERN2 may try to access a port that is already being used - if the process using the port is not important, it can be killed using the following command `fuser -k [port number]/tcp`. If BERN2 is not running properly, it can be restarted cleanly by: deleting the “log/” folder, stopping BERN2 using the appropriate script, and deleting any CUDA artefacts.

Once the BERN2 model is running on a local port within its conda environment, we make requests to it using the URL associated with the port. Running our corpus through the model returns a set of found entities per paper (and optionally per paragraph) which fall into one of nine biomedical classes: gene/protein, disease, drug/chemical, species, mutation, cell line, cell type, DNA, and RNA. We filter entities for the gene/protein class to enable protein NER comparison to our model.

### Applicability of our model to improve metabolite NER

The metabolite NER (32) and enzyme NER tasks were carried out independently, and hence use different training, validation and test sets. To compare the output of both algorithms we evaluated the overlap of the two test sets, yielding a total of 18 metabolomics publications that have not been used as part of the training or validation stages in either algorithm. We extracted all sentences with manually annotated metabolites and/or enzymes and compare the outputs of both NER algorithms to evaluate whether our model can help to improve the accuracy of the metabolite NER output.

### Evaluation Metrics

To evaluate the performance of the annotation pipeline and DL models, we calculate 3 properties of a confusion matrix: true positive (TP) entities, false positives (FPs) and false negatives (TNs). FPs are entities predicted to be an enzyme by a model which does not match an entity in the test set. FNs are entities that were not annotated by the model but are contained in the test set annotations. Using these measures, we calculate the precision (probability of correctly predicted positive values in the total predicted positive outcomes, Equation 1), recall (also known as sensitivity, probability of correctly predicted positives relative to all true positive cases, Equation 2) and F1-score (weighted average of precision and recall, Equation 3).

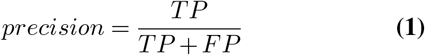

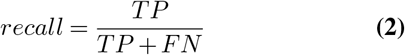

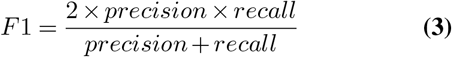

Partial positive (PP) matches, i.e. where the only root term is correctly identified or where the indication of an isoform is missing but the remainder of an entity is correctly annotated, were separately tracked and the final count of PPs was split equally between TP and FN.

## Results and Discussion

### Corpus creation and hierarchical dictionary

The reconstructed dictionary list, using the hierarchical format (Figure 1), speeds up the searching process compared to a standard dictionary search. An entire full-text article can be annotated in 0.5s using this approach, compared to 1s per file using the standard approach, due to the reduction in size of the branch list resulting in less evaluations.

**Fig. 1.**
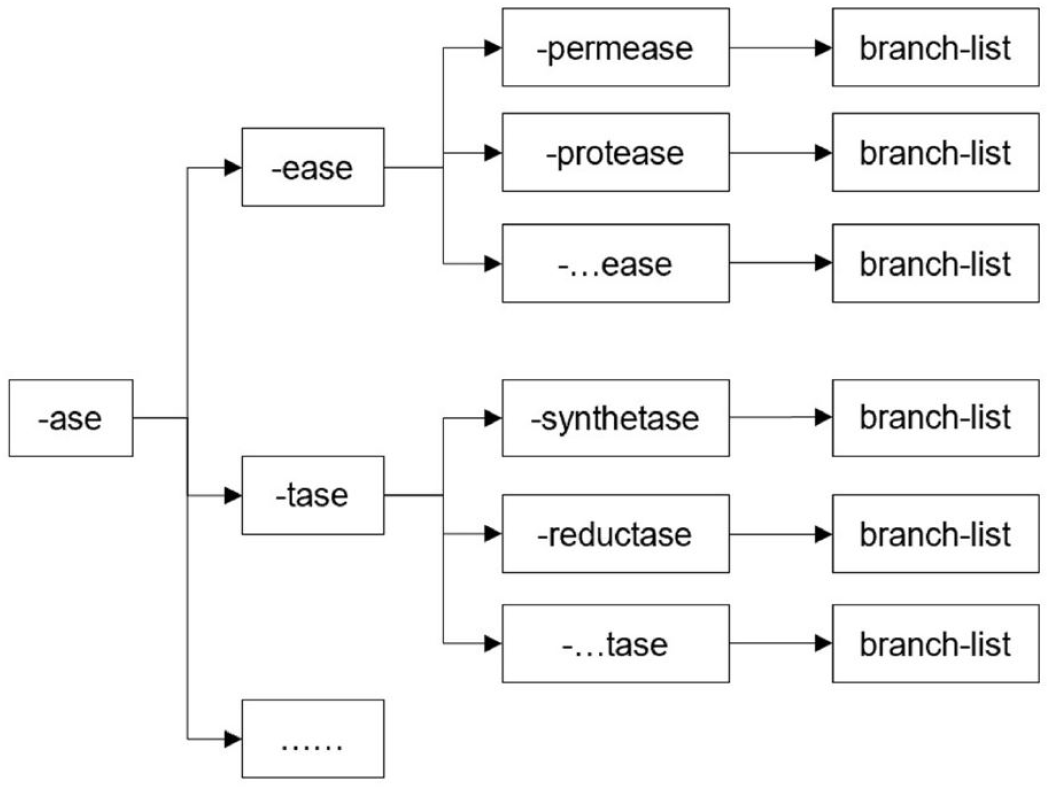
Example of the hierarchical structure of the dictionary showing a part of the ‘-ase’ branch. Enzymes not ending in ‘-ase’ are contained in the dictionary in other branches.

Our annotation pipeline was used to create an annotated corpus for supervised learning. The enzyme entities in the texts are extracted and coded into the “annotations” section of the BioC JSON file (output of Auto-CORPus) that follows the text content of each paragraph, see example in Figure 2. Each annotation has a number of key-value pairs: ‘text’ is the name of the identified entity, ‘infons’ contains the information on the identifier (E.C. number, or keyword), type (enzyme), annotator, annotation time, ‘id’ is a unique ID for each annotation in the text and indicates the order in which the entity is annotated, and ‘locations’ indicates the position of the annotation within the full-text.

**Fig. 2.**
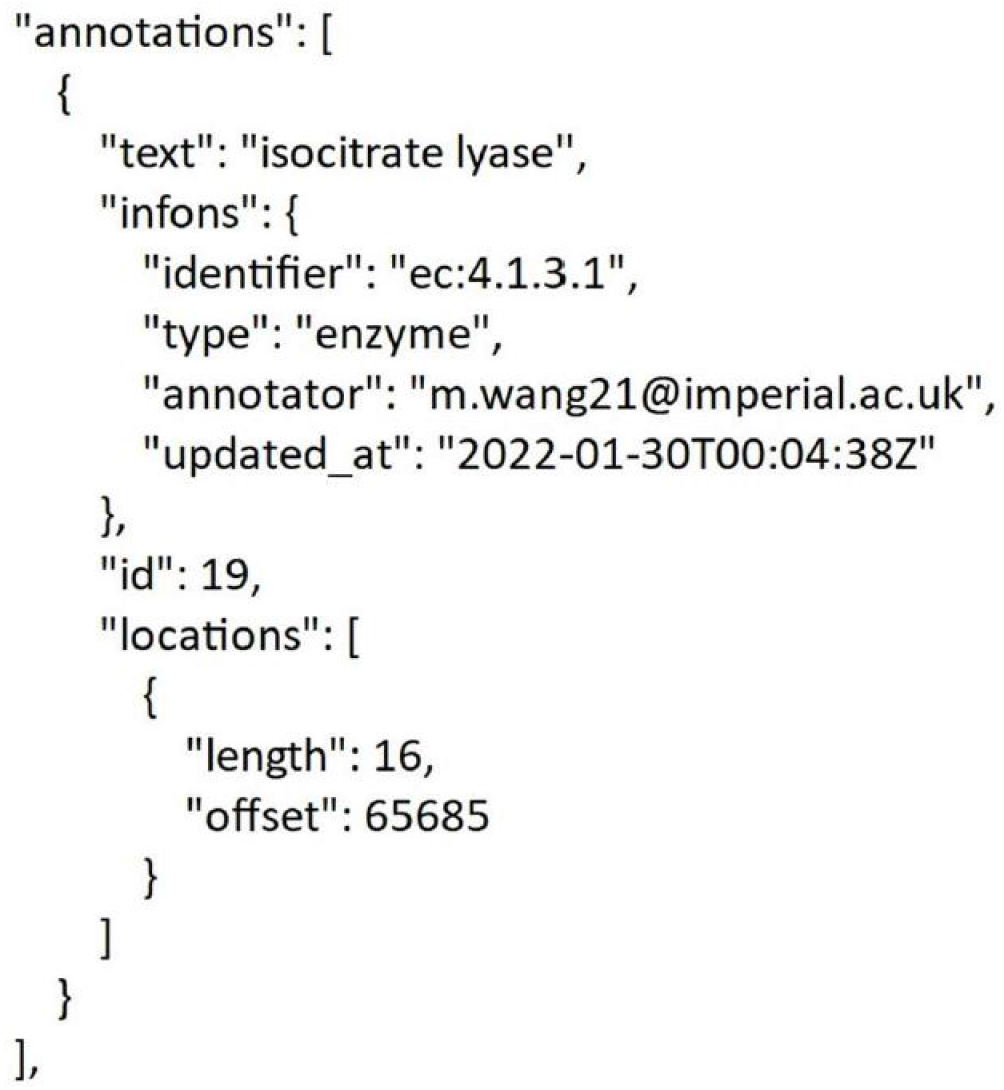
An example of annotations. That is an individual dictionary that displays the detail of an entity, which has four elements (i.e., ‘text’, ‘infons’, ‘id’, ‘locations’) to represent a unique entity in the text.

The annotated corpus is easy to use, and can be reused in future to convert the text and annotations to different formats to be used for training different algorithms. Moreover, the hierarchical annotation approach may also be applicable for speeding up the process of annotating other biomedical entities such as pathways, gene names or metabolites. For example, pathways also have a similar nomenclature as enzymes in which entities end in similar ways such as the root-words ‘synthesis’, ‘biosynthesis’, ‘degradation’ and ‘metabolism’. The JSON structure also lends itself to contain multiple entity types which means multi-task learning corpora can be prepared for biomedical NER tasks.

However, our pipeline includes some modifications to improve the accuracy and effectiveness of annotation. The weakness of the annotation pipeline mainly lies in the second searching step (i.e., keyword searching). Once a root word has been identified, the preceding and following words are evaluated whether these are part of the enzyme. The dictionary contains several spellings of an enzyme, however different forms may still exist that may include (or exclude) the use of hyphens which will prevent terms to be matched directly. Likewise, different isoforms may be indicates by added numbers or letters at the end. Our annotation pipeline uses a rule-based regular expression search to evaluate if the next word may indicate an isoform, however this approach is not fool-proof. Finally, another difficulty is that ambiguous semantics and multiple expressions of enzymes are unfavourable for annotators to precisely distinguish an enzyme name within the sentence. For example, some descriptive modifiers such as ‘human’, ‘serum’, or ‘bacterial’ are truly parts of an enzyme name, like ‘human carboxylesterase 2’ (EC:3.3.3.84), but more generally, these words should be simply regarded as the adjectives of enzymes, such as ‘human proteases’ and are therefore not included in our annotations unless for some cases where there are exact matches. The difficulty of manual annotation highlights how this process may be replaced by an algorithm before being subjected to human verification.

Our training data set created from the annotated JSON files additionally replaces all abbreviations of enzymes by their definitions, hence the evaluation of abbreviated entities relies on correctly identifying abbreviation-definition pairs. Our annotation pipeline annotated over 30,000 entities in over 3,000 articles (Table 2). As expected, the proteomics articles have the largest average number of entities (23) followed by metabolomics (9) and GWAS (8). Across all annotated entities, the most common single word annotation is polymerase, whereas when including multi-word entities the most common root terms are kinase and protease (Table 3). No enzyme-specific corpus exists, and our corpus is also larger than most bioNLP corpora (38).

**Table 2.** Summary of the annotated enzyme corpus.

| Domain | Annotated enzymes | Mean | St. dev. |
| --- | --- | --- | --- |
| GWAS | 6,736 | 8 | 17.77 |
| proteomics | 10,037 | 23 | 29.30 |
| metabolomics | 9,447 | 9 | 13.76 |
| microbiome | 5,502 | 5 | 7.88 |
| total | 31,722 | 9 | 17.11 |

**Table 3.**
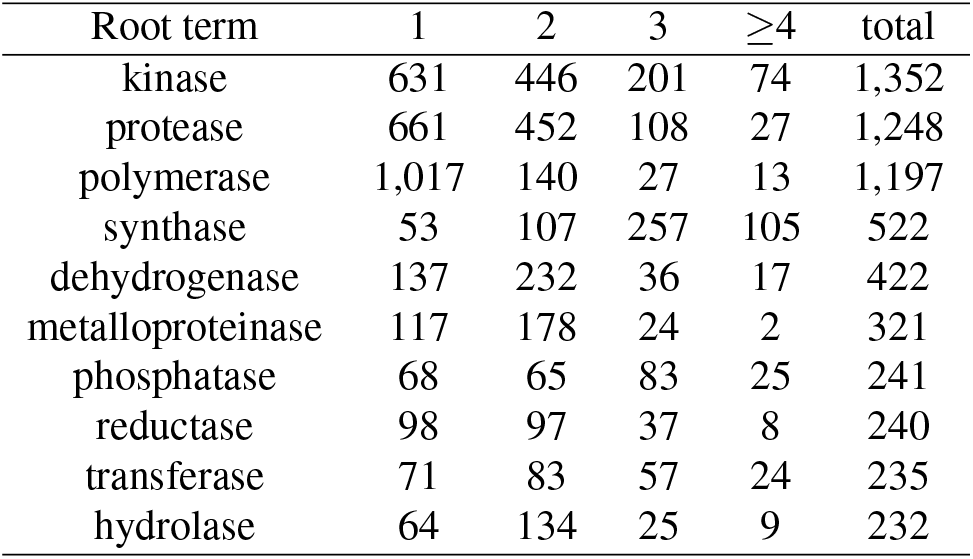
The number of components of entities identified by keyword searching for the 10 most found root terms separated out by the number of words.

### Model Evaluation

As described in the results, we tracked all partial matches and split these between TPs and FNs. We also analysed the list of PP items. Some of these PPs are similar (but not identical) between the manually annotated test set and the annotation (or DL) pipeline. The main differences related to a capital letter or a number not captured by the annotation pipeline (i.e. isoforms). Most other differences relate to the method only identifying the root term (‘kinase’ instead of ‘Ser/Thr kinase’). The former being a minor mistake (could be considered a TP), whereas the latter clearly misses an important part of the enzyme (and perhaps should be a FN). Resolving PPs into TPs/FNs would involve a fourth independent person (from the annotators and arbiter) to go over each of these entries which was not feasible given the time/effort required. Therefore here we regarded half of the PPs as TP and the other as FN.

The precision of the pipeline is effectively 1.00 (see Table 4), and there is only one false positive (‘protein A’). However, the lower recall of 0.69 demonstrates that there are certain enzyme entities that were not found by the annotation pipeline (higher degree of FNs). Most of these missed entities are abbreviations (with added characters or incomplete matches) which is hard to identify without the understanding of the context in which entities such as ‘CN-II’ or ‘dN-1’ first appear, for this reason we separately evaluated the results in Table 4 while omitting abbreviations from the calculations, which resulted in a recall of 0.76. A logical next step is to further improve abbreviation detection algorithms (currently using rule-based methods) by taking a DL approach to focus on relation extraction between abbreviations and definitions. Another aspect we identified is that punctuation and hyphenation also impact the performance of the annotation pipeline (e.g. ‘dipeptidyl peptidase-4’ and ‘coenzyme A-linked enzyme’ were not identified by the pipeline). Further improving the pipeline requires both programming new rule-based algorithms and a human-in-the-loop system to more effectively devise rules to improve the accuracy of the automatic annotation pipeline.

**Table 4.** Performance metrics for different models evaluated on the manually annotated test set (n = 526 full text articles).

| Model | Embedding | Evaluation | Precision | Recall | F1-score |
| --- | --- | --- | --- | --- | --- |
| Annotation pipeline | - | All | **0.995** | 0.688 | 0.813 |
| BioBERT-BiLSTM | BioBERT (12) | All | 0.968 | 0.940 | 0.950 |
| SciBERT-BiLSTM | SciBERT (11) | All | 0.966 | **0.951** | **0.956** |
| BERN2 (15) | BioBERT (12) | All | 0.134 | 0.856 | 0.231 |
| Annotation pipeline | - | Excl. abbreviations | **0.996** | 0.762 | 0.863 |
| BioBERT-BiLSTM | BioBERT (12) | Excl. abbreviations | 0.981 | 0.937 | 0.955 |
| SciBERT-BiLSTM | SciBERT (11) | Excl. abbreviations | 0.981 | **0.954** | **0.965** |
| BERN2 (15) | BioBERT (12) | Excl. abbreviations | 0.148 | 0.824 | 0.250 |

As an alternative to dictionary searching combined with rule-based annotation we created two DL models based on the BiLSTM architecture that achieved state-of-the-art results in metabolite named-entity recognition (32). Here we extended that work, in the context of enzyme NER, by comparing the best embedding found previously (BioBERT (12)) with the newer SciBERT (11) model which includes its own vocabulary. Both methods achieve very similar precision (0.97) on the full test data, which increases to 0.98 when excluding abbreviations, however their recall far exceeds the annotation pipeline with 0.94 for the BioBERT-BiLSTM and 0.95 for the SciBERT model. This is also reflected in the F1-scores of 0.96 for SciBERT-BiLSTM, 0.95 for BioBERT-BiLSTM and 0.81 for the annotation pipeline. When excluding abbreviations these increase to 0.97 (SciBERT-BiLSTM), 0.96 (BioBERT-BiLSTM) and 0.86 (annotation pipelines). This highlights the benefit of using DL algorithms for NER over (traditional) dictionary-based approaches where a slightly lower precision for DL methods is accompanied with a substantially higher recall.

Intuitively, we would expect that the BioBERT-based model would yield better performance than SciBERT, since it was specifically pretrained on a biomedical corpus. However, we find that the SciBERT-based model (for tokenisation and work embedding) has the best performance overall. SciBERT (11) was trained on a corpus that contains over 80% of papers from the broad biomedical domain which results in the transformer learning features of the words frequently used in biomedical literature. In addition, SciBERT constructed a new vocabulary, SCIVOCAB, for tokenisation and word embedding which has a substantial difference from the Word-Piece vocabulary used by BERT and inherited by BioBERT (which was built on top of BERT). As Beltagy et al. indicate, the overlap between the two vocabularies of tokens is a mere 42%, and SCIVOCAB provides more detailed tokens with biomedical features (11).

With the development of biomedical NLP, more and more DL architectures are built or pretrained as transformer models, with BioBERT (12) being the most popular. BioBERT has been reused by the same authors to create a model (BERN2) for joint NER and named entity normalisation for 9 different entities including proteins (achieving an F1-score of 0.84 on the BC2GM corpus (24)). To the best of our knowledge, our work is the first time in which algorithms are developed for an annotation pipeline and NER models specifically for enzymes (opposed to all proteins). We therefore evaluate the performance of the BERN2 model on our manually annotated enzyme corpus and compare it with our enzyme-specific models. Since not all proteins are enzymes, and hence there can be many false positives, we evaluate BERN2 using the recall only. BERN2 achieves a recall of 0.86, clearly out-performing our dictionary-based annotation pipeline but not reaching the same level as our dedicated enzyme NER models (recall 0.94-0.95).

It remains to be seen whether a pure transformer model is able to reach similar results and this would form a logical next step of the work that can build on our creation of a corpus for enzyme-specific NER. Moreover, our annotated data contains over 30,000 entities (Table 2) compared with approximately 20,000 for the BC2GM corpus (24). While the protein NER corpora will include enzymes in the training data, our training set is much larger which means that the models can be trained and evaluated with more data. Moreover, the methods used here for enzymes are amenable to any type of protein to facilitate the creation of a multi-task protein NER model and other biomedical text-mining tools.

The evaluation of the NER tasks is a significant step of improving the training model performance. In this study, our metrics for the DL models were generated based on tokens, i.e., the model classifies every token to its unique label from the (‘S’, ‘O’, ‘B’, ‘I’, ‘E’) scheme, and we have a classification evaluation that calculates these terms of TP, FP, FN and then compute the precision, recall, and F1-score. This evaluation was used to get the exact accuracy for each token, but there are other, potentially more realistic, schemas to evaluate these models that consider partial matches such as the PPs. The Message Understanding Conference (MUC) (39) has introduced some metrics for evaluation considering the different categories of misclassified values against the strict classification, with the International Workshop on Semantic Evaluation (SemEval) resulting in the measurement to calculate corresponding precision, recall, and F1-score based on the MUC metrics. However, while these metrics give a better representation of the errors and performance, they are not used in the evaluation of modern NER models and as such would not make it possible for us to compare our results with models such as BERN2.

### Model Application to Improve Metabolite NER

Our prior work (32) showed that some entities recognised as metabolites are actually part of enzyme names and hence ought to be considered as false positives when evaluating these in the context of metabolite NER. Our work here on enzyme NER can be used to address this limitation by filtering out any metabolite entities that overlaps with an enzyme entity.

We show in Table 5 several examples of applying both TABoLiSTM and SciBERT-BiLSTM to the same sentence. We have repurposed the TABoLiSTM metabolomics corpus (over 1,000 publications in that corpus contain at least 1 enzyme, see Table 1) in this work, therefore in order to evaluate how SciBERT-BiLSTM can improve TABoLiSTM we only give examples from articles that were part of either test set. Table 5A contains examples where we identified no benefit of using SciBERT-BiLSTM to improve the output of TABoLiSTM as no entities overlap, hence TABoLiSTM correctly did not identify ‘tyrosine’ as metabolite as it was part of an enzyme name (it was not trained with this purpose). In Table 5B there are several examples where TABoLiSTM reports a metabolite that is actually part of an enzyme (SciBERT-BiLSTM output) such as identifying ‘ceramide’ twice in a sentence where only one of these is a metabolite and the other is part of ‘ceramide synthases’, or identifying ‘deoxycytidine’ as metabolite when SciBERT-BiLSTM identifies it as ‘deoxycytidine kinase’. Moreover, TABoLiSTM (32) may also have an application to our enzyme NER work here as it can be used to identify whether there is a metabolite name directly in front of a root term (in cases where only a root term is identified). This demonstrates that these models can be used together to improve each other, while this can be done in the context of employing these algorithms in parallel, other work (15) has shown that multi-task learning will yield both better performance and small model sizes.

**Table 5.** Example output from metabolite NER (32) (underlined) and enzyme NER (in bold). Correct annotations are indicated in green, false positive metabolite annotations within enzymes in red, and false negative metabolite annotations in orange. For purposes of diversity of metabolites and enzymes we show one example per article to illustrate different cases. Additional examples are in the Supporting Information in Table 6.

| A. No metabolites annotated within enzymes |
| --- |
| PMC4525767: “However, another equally plausible hypothesis is that reductions in the activity of **branched chain ketoacid dehydrogenase** (BCKD) and **tyrosine aminotransferase** (TAT) in states of insulin resistance lead to increased tissue and circulating BCAA and AAA concentrations, respectively [19].” |
| PMC6316856: “Serum <u>glucose</u>, HbA1c, <u>triglycerides</u>, total <u>cholesterol</u>, <u>LDL</u> and <u>HDL-cholesterol</u>, <u>alanine aminotransferase</u> (ALT), <u>aspartate aminotransferase</u> (AST), <u>gamma-glutamyl transpeptidase</u> ($\gamma$ GT), <u>creatinine</u>, <u>uric acid</u>, <u>vitamin D</u>, <u>folic acid</u>, ferritin, c-reactive protein, <u>thyroxin</u> and thyroid-stimulating hormone were measured in a certified clinical laboratory, using standard protocols.” |
| PMC6875299: “In the <u>glucose-alanine cycle</u> pathway, <u>glutamate dehydrogenase</u> in muscle catalyzes the binding of <u>α-ketoglutaric acid</u> to <u>ammonia</u> to form <u>glutamate</u>, followed by <u>glutamate</u> catalyzed by <u>alanine aminotransferase</u>; <u>pyruvic acid</u> forms <u>alpha-ketoglutarate</u> and <u>alanine</u> [30].” |
| B. Annotation of metabolite within an enzyme, false positive metabolite annotations can be filtered out |
| PMC3406255: “The 4-5 doublet in <u>glutamate</u> and <u>glutamine</u> carbon 4 is derived from <u>[1,2-13C]acetyl-CoA</u> produced from <u>[U-13C]glucose</u>, demonstrating that <u>glucose</u> was metabolized to <u>acetyl-CoA</u> via <u>pyruvate dehydrogenase</u> (PDH).” |
| PMC3645730: “In fact, higher levels of <u>ceramide synthases</u> and <u>ceramide</u> were detected in breast cancer tissues compared with those in adjacent normal ones [42].” |
| PMC5073280: “Undiluted serum sample (25 μL) and 100 μL of 2 mM <u>hydrogen peroxide</u> was added to freshly prepared reaction mixture consisting of 755 μL of 0.1 M <u>sodium phosphate</u> buffer (pH 7.0), 100 μL of 2 mM <u>nicotinamide adenine dinucleotide phosphate</u> (NADPH), 10 μL <u>glutathione reductase</u> (2 μL in 500 μL buffer) and 10 μL reduced <u>glutathione</u> (9.22 mg/mL) kept in cuvette of 1 mL capacity.” |
| PMC5287439: “As a membrane constituent, <u>sphingomyelins</u> are implicated in transmembrane signaling and are generated from <u>phosphatidylcholine</u> and <u>ceramide</u> by <u>sphingomyelin synthase</u>, the knockout of which leads to mitochondrial dysfunction and reduced insulin release.” |
| PMC6224486: “MG00/1846Z,9Z,12Z,15Z/00, another metabolite in the panel, is a <u>monoacylglyceride</u> that can be broken down by <u>monoacylglycerol lipase</u>.” |
| PMC6708594: “Once inside the cell, <u>fludarabine</u> is sequentially phosphorylated to the <u>monophosphate</u> (F-ara-AMP), <u>diphosphate</u> (F-ara-ADP), and <u>triphosphate</u> (F-ara-ATP) forms by <u>deoxycytidine kinase</u> (dCK), <u>adenylate kinase</u> (AK), and <u>nucleoside diphosphate kinase</u> (NDK), respectively [32].” |

**Table 6.**
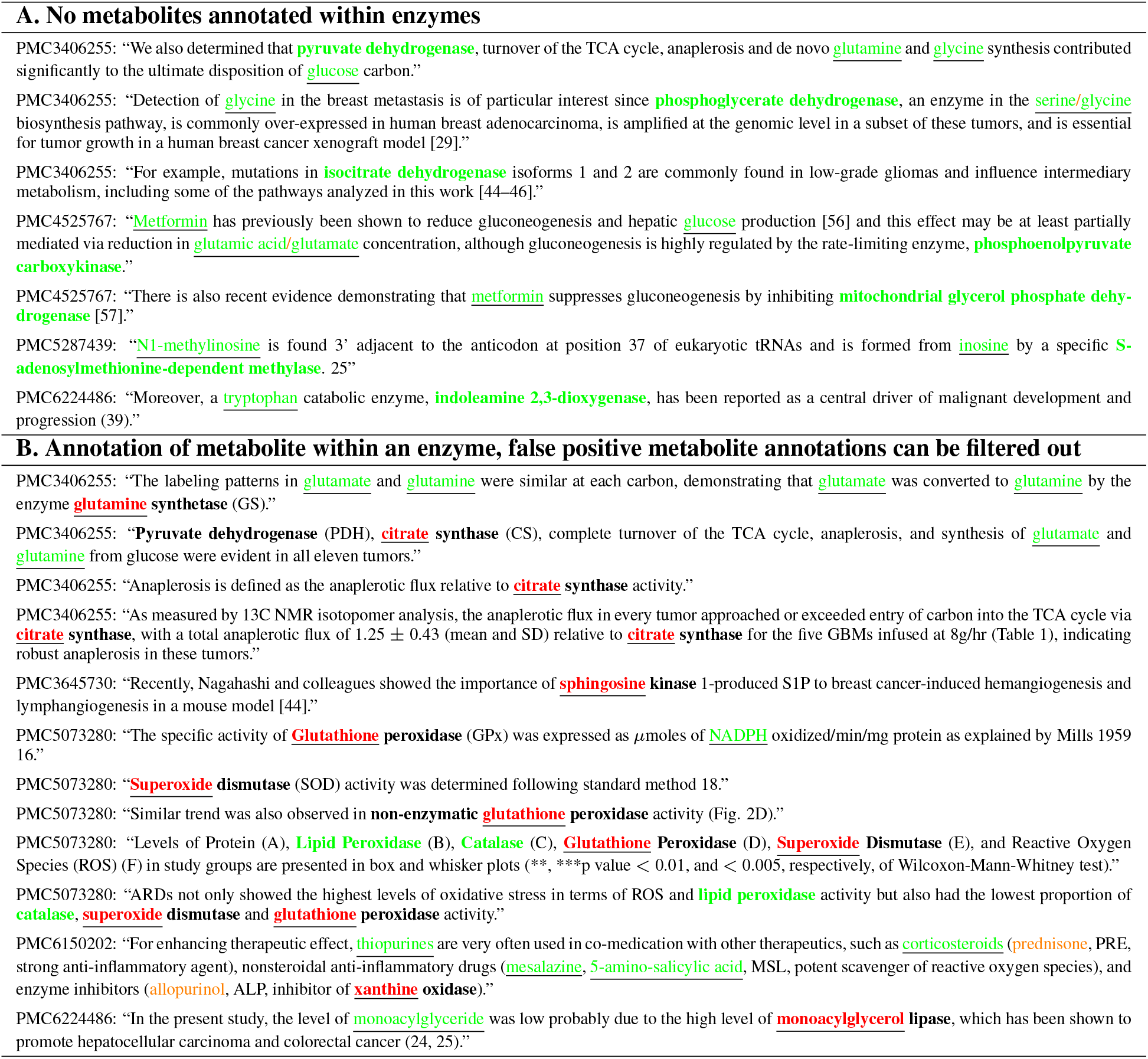
Example output from metabolite NER (32) (underlined) and enzyme NER (in bold). Remaining set of sentences (in addition to Table 5) for all 18 articles used for the evaluation (see Materials and Methods). Correct annotations are indicated in green, false positive metabolite annotations within enzymes in red, and false negative and other false positive metabolite annotations in orange. Note: formatting (e.g., superscript) is removed after processing with Auto-CORPus (31) before NER.

## Conclusions

In conclusion, this work contributed a novel dictionary- and rule-based automatic pipeline for enzyme NER with almost perfect precision and which runs in a matter of seconds (on a standard laptop) on an entire full-text article because of the hierarchical dictionary. We used this pipeline to create the first-of-its-kind corpus for enzyme NER to facilitate text-mining. Our best performing model was the SciBERT word embedding and tokenisation combined with a BiLSTM model and this deep learning model achieved a state-of-theart performance (F1-score*>*0.95) for enzyme NER. Our proposed pipeline with the DL models can improve more effective enzyme text-mining and information extraction research for literature review and be used to improve other biomedical NER tasks.

The code for automatic annotation, model training and inference, as well as the data for training and testing are available at https://github.com/omicsNLP/enzymeNER.

## AUTHOR CONTRIBUTIONS

Conceptualization, J.M.P. and T.B.; Methodology, M.W., J.M.P. and T.B.; Formal Analysis and Investigation, M.W., A.V. and J.M.P.; Writing – Original Draft: M.W.; Writing – Review Editing: J.M.P., M.W., T.B and A.V. All authors have read and agreed to the published version of the manuscript.

The authors thank Cheng Su Yeung and Nazanin Faghih Mirzaei for their support and help during the code development.

A.V. is supported by a UK Research and Innovation (UKRI) Centre for Doctoral Training in AI for Healthcare PhD studentship (EP/S023283/1 (EPSRC)). This research was funded by Health Data Research (HDR) UK and the Medical Research Council (MRC) via an UKRI Innovation Fellowship to T.B. (MR/S003703/1) and a Rutherford Fund Fellowship to J.M.P. (MR/S004033/1). The authors declare no conflict of interest. The funders had no role in the design of the study; in the collection, analyses, or interpretation of data; in the writing of the manuscript, or in the decision to publish the results.

## Notes

https://github.com/omicsNLP/enzymeNER

### Competing Interest Statement

The authors have declared no competing interest.

